# Unexpectedly dense colonization of soil pore space by microbes

**DOI:** 10.1101/2024.04.10.588810

**Authors:** Hannes Schmidt, Steffen Schlüter, Xavier Raynaud, Vincent J.M.N.L. Felde, Berit Zeller-Plumhoff, Andreas Richter, Naoise Nunan

## Abstract

Pore surfaces in soil are considered to be sparsely colonised, dispersed only with isolated cells or colonies of bacteria and archaea. Here, we question this ‘empty space’-concept by combining microstructure analysis with molecular biology and provide a data-driven update on habitable surface areas (HSA) in soil. Our unique approach allowed us to provide 1) evidence that microbial densities in soil have been underestimated for decades and 2) two-dimensional simulations of their potential distribution within the soil pore space. Our results demonstrate the need for a new perspective on how densely soil is colonised, with implications for how we think of basic ecological processes such as microbial motility or predation, and how microbial processes, including organic matter dynamics, are ultimately modelled.

## Main text

Soil is a complex habitat that features a multitude of ecological niches. This complexity is considered to be the main reason for the vast microbial diversity, not only on a global, but also on a micrometer scale [1,2]. Each gram of soil contains 10^8^ - 10^10^ bacteria and archaea and provides internal surfaces for microbial colonization within pore space. Soil microbes are regarded as the main engines of biogeochemical cycling and there is evidence to suggest that microbial community functioning requires a certain density of cells to allow for cell-cell communication, gene expression and regulation of energy-costly processes such as enzyme production or motility [3–7]. However, soil bacteria and archaea are believed to be isolated in soil pore space [8–13]. Young et al. [8] estimated that the total pore surface area covered by soil protozoa, fungi, and bacteria is in the order of 10^-6^ %, leading to the idea that cell-cell distances were large, with an average of up to 500 μm between individual cells or microcolonies [1].

In this brief communication we present evidence that the density of bacteria and archaea in soil is orders of magnitude higher than previously estimated. We used aggregates (1-3 mm) from a temperate forest topsoil to analyse (i) the HSA via X-ray microtomography at the relevant resolution (2 μm) to resolve most of the habitable pores as well as (ii) the abundance of bacteria and archaea via droplet digital PCR. Combining these data allowed us to calculate the proportion of HSA covered by cells and to simulate the distribution and density of microbial cells in soil pore space.

X-ray microtomography revealed a surface area density of 14.6 mm^2^/mm^3^ which translated to a HSA ranging from 64.2 - 168.1 mm^2^ and microbial numbers ranging from 1.9 - 9.3 × 10^6^ cells for four aggregates (Table 1; Data S1). Assuming a random distribution, the distance to the nearest neighbouring cell varied from 1.6 to 3.8 μm and the number of neighbouring cells within an interaction distance of 20 μm ranged from 29 to 178. A similar number of bacterial neighbours in soil has been suggested before [14], albeit not within the context of soil pore space where the cell-cell distances may even be smaller when pores are saturated with water. Our data suggests a coverage of the available habitable surface area of 1.0 to 5.8 % by microorganisms with an average cell size of 0.75 μm (Table 1). These calculations indicate that soil surfaces are much more heavily colonised by bacteria and archaea alone than the currently accepted value for bacteria, archaea, fungi, and protists altogether [8]. Previous estimations were only based on theoretical values and often used surface areas of clay minerals based on gas adsorption analysis, thus probing at a scale of molecules and not at the scale of microbial cells. Surfaces of clay platelets contribute massively to gas adsorption but are too small to fit individual bacteria. Consequently, previous studies overestimated the HSA while often underestimating microbial abundance (e.g. [8]). In contrast, the use of high-resolution X-ray microtomography allowed us to assess HSA more accurately, as it measures the actual surface area at the scale of cells and takes the spatial arrangement of particles into account (Figure S1). Combined with microbial abundance analysis, these data enabled us to show that bacterial and archaeal densities in soil are unexpectedly high.

**Table 1.** Densities of microbial cells with an average diameter of 0.75 μm per habitable surface area of four individual aggregates obtained via X-ray microtomography and droplet digital PCR.

| Aggregate | HSA per<br>aggregate [$\text{mm}^2$] | Cells per<br>aggregate [n] | Cells per HSA<br>[n/10,000 $\mu\text{m}^2$] | Neighbouring<br>cells [n] | HSA covered<br>by cells [%] |
| --- | --- | --- | --- | --- | --- |
| A1m | 64.2 | 3.7E+06 | 577 | 78 | 2.5 |
| A2m | 70.6 | 9.3E+06 | 1313 | 178 | 5.8 |
| A3m | 168.1 | 8.3E+06 | 493 | 67 | 2.2 |
| A4m | 87.9 | 1.9E+06 | 216 | 29 | 1.0 |

Bacteria and archaea are most likely not randomly distributed in soil pore space. We used a few simple rules supported by microscopic observations to illustrate the potential distribution of cells in a two-dimensional space representing the surface of soil meso- and macropores (Figure S2 and Supplementary Material). These simulations show a microbial coverage of 2.8 % of HSA on average (Figure 1A). Smaller subplots reveal local colonization heterogeneity (Figure 1B and C), suggesting varying interaction strengths within soil. Community composition showed marked differences at the microscale and spatial variability was pronounced for the less abundant phyla such as Actinobacteriota or Chloroflexi (Figure 1) but was also visible for the dominating group (Proteobacteria) under low and high colonization densities (Figures S3 and S4). Plotting the densities of bacteria and archaea per HSA for each individual aggregate revealed large differences in their potential to interact with neighbouring cells (Figure S3 and S4), indicating that this specific forest soil hosts microbial communities of variable composition at the microscale. As a result, community assembly and priority effects could influence local processes performed by soil bacteria and archaea [15–17].

**Figure 1.**
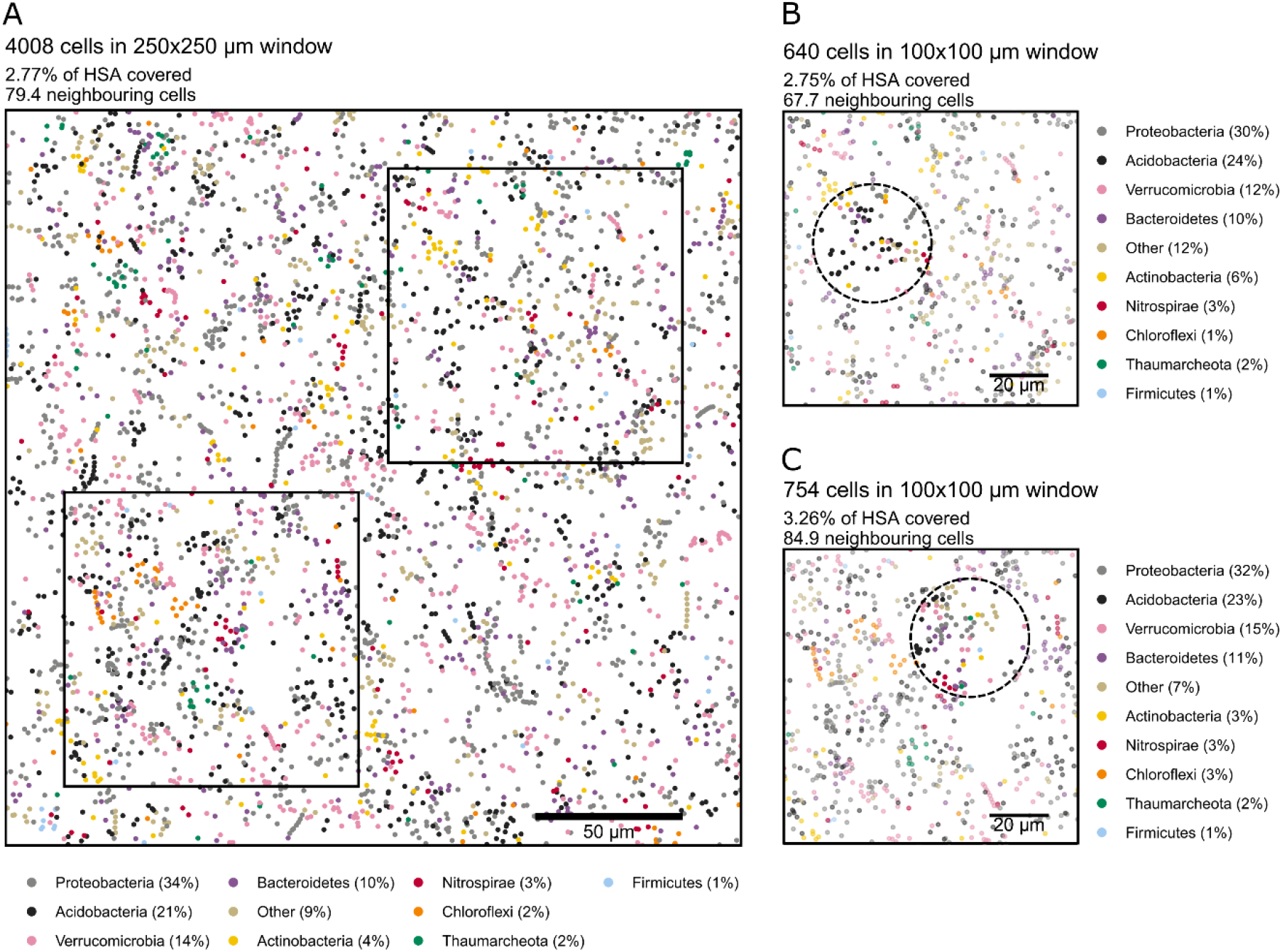
Two-dimensional simulation of bacterial and archaeal colonization of habitable surface area in soil. Panel A shows a simulation of microbial cell density (cell size 0.75 μm) based on the average cell number and community composition for four aggregates plotted on an area of 0.0625 mm^2^ (250x250 μm window). The average coverage of HSA was 2.8 %. At this density, each cell had approx. 80 neighbouring cells within an interaction distance of 20 μm. Panels B and C are subsets of panel A and show the potential interaction distance of 20 μm (black circle) around a single cell. Note that cell numbers, HSA covered, neighbouring cells, as well as the relative abundance of individual groups showed marked differences between subpanels indicating heterogeneous distribution of microbial taxa within their soil microenvironment. Colours were randomly assigned to bacterial and archaeal groups and cells were distributed as micro-colonies (55 %), single cells (40 %), or filaments (5 %). Cell abundance and community composition was assessed via quantitative microbiome profiling and surface area was obtained via X-ray microtomography (see Supplementary Material for details). Scale bars: A: 50 μm; B,C: 20 μm.

To test if our findings were transferrable to other soils, and to see whether the scanning resolution in this study (2 μm) potentially underestimated the HSA, we established a power-law model fit to 18 datasets that resulted in an average soil surface density of 161.8 mm^2^/mm^3^ at a scanning resolution of 1 μm (Supplementary Material; Data S2; Fig. S6). In addition, we established an average bacterial and archaeal cell number of 3.2 × 10^9^ cells g^-1^ soil obtained from 13 datasets (Data S2). Combining these data allowed us to estimate the average proportion of HSA covered by microorganisms to be 0.3%, 1.1 %, and 2.8 % for cell sizes of 0.5 μm, 0.75 μm, and 1 μm, respectively (Data S2). These values are highly congruent with our experimental data above (Table 1) and support our findings that the local density of microbial cells per HSA is magnitudes higher than previously estimated (Figure S5). This calls for a general revision of microbial densities in soil.

## Acknowledgments

We would like to thank Sean Darcy, Christina Kaiser, as well as Kobler Johannes and the IM LTER Zöbelboden Team for assistance in field work and meta data preparation. We further thank Leila Jensen for excellence assistance in the laboratory. We thank Petra Pjevac and Bela Hausmann of the Joint Microbiome Facility of the Medical University of Vienna and the University of Vienna, Austria, for assistance with amplicon sequencing and data pre-processing. We specifically thank Joana Senéca Silva for handling the submission of the sequencing data to the NCBI Short-Read Archive. For open access purposes, the author has applied a CC BY public copyright license to any author accepted manuscript version arising from this submission.

## Author contributions

HS conceived the study in consultation with NN. Data analysis and processing was done by BZ-P (microtomography, scanning), SS (microtomography, data analysis), VF (microtomography, data analysis), XR (bacterial simulations) in consultation with HS who synthesised the data. HS wrote the manuscript with input from all co-authors.

